# A new locally collected environmental quality indicator for medium and large mammals

**DOI:** 10.1101/2024.06.10.598337

**Authors:** Márcio Leite Oliveira, Guilherme Rossi Gorni, Alessandra Santos Nascimento, Fernando de Camargo Passos

## Abstract

Measuring environmental degradation with bioindicators, landscape metrics, and remote sensing helps understand impact on biota. However, data on anthropogenic pressures such as plant exploitation, poaching and invasive species are crucial. We created an Anthropogenic Influence Index (AII) for medium and large mammals at the Atlantic Forest based on local environmental quality indicators and tested its correlation with existing indices, such as the Global Human Influence Index (GHII), landscape metrics and social-economic indicators. We found no correlation between the AII and the GHII, indicating that remote sensing-collected data may not reflect local and specific anthropogenic impacts on the environment. In addition, there was a correlation between the AII and the Human Development Index, drawing attention to the direct relationship between income, education and life expectancy and the incidence of environmental impacts. Thus, the AII appears to better capture local nuances of environmental impacts, particularly those significant for medium and large mammals, compared to other indicators such as GHII, human density, and landscape metrics.

## INTRODUCTION

The surge in human population drives increased resource consumption, expanding the human footprint on the environment. This interaction between population growth and resource usage has varying ecological impacts, shaped by local contexts and species involved. Furthermore, the link between environmental changes and development hinges on each country’s energy and production infrastructure, affecting resource utilization and environmental management. Additionally, historical human activity in diverse regions profoundly shapes ecosystem outcomes (Sanderson et al. 2002).

Measuring the level of environmental degradation in habitats due to human action is important for understanding how degradation impacts biota, monitoring its influence over time, and ultimately understanding how human action affects ecosystems. These measurements have been carried out in several ways, such as using bioindicators (Holt and Miller 2010) landscape ecology metrics (Uuemaa et al. 2013) and remote sensing combined with human influence indices (Wildlife Conservation Society and Center for International Earth Science Information Network 2005). Thus, some variables serve as proxies for environmental degradation, such as human population density (Spear et al. 2013), purchase power parity (Nordhaus 2006), and the Humam development index (Jha and Bawa 2006).

A widespread used index of anthropogenic influence level on the environment is the Global Human Influence Index (Wildlife Conservation Society and Center for International Earth Science Information Network 2005). It is a global dataset of 1-kilometer grid cells, composed of nine global data layers covering human population pressure (population density), human land use and infrastructure (built-up areas, nighttime lights, land use/land cover), and human access (coastlines, roads, railroads, navigable rivers). It has been used to indicate conservation priority areas (Oliveira et al. 2022), to evaluate human impact on avian distribution (Yohe et al. 2014), as proxy of propagule pressure of alien species (Gallardo et al. 2015), hunting pressure and is related to ungulates density (Martínez□Gutiérrez et al. 2018). However, it may not be adequate for some situations once it may fail to incorporate local nuances.

Tropical and subtropical moist broadleaf forests (Olson et al. 2001) are the most diverse terrestrial environments, harboring half of earth’s species (Groombridge and Jenkins 2002). These are biomes worldwide under several threats such as at the Amazon region (Silva Junior et al. 2020) and the Asia Southeastern (Stibig et al. 2014). Examples of threats are the expansion of cultivated areas (Rajão et al. 2020), introductions of invasive species (Hegel et al. 2019), poaching (Bogoni et al. 2020) and predation by domestic animals (Cassano et al. 2014). One of these forests are the Atlantic Forest that is spread over the Brazilian southern region and eastern coastline, the eastern Paraguay and the Missiones province in the Argentina, holding 746 vertebrate species (Mittermeier et al. 2011). It has lost 88% of its original cover (Ribeiro et al. 2009a) and has a high human population density (Cincotta et al. 2000). At the Atlantic Forest, as in other rainforests, medium and large mammal species are particularly threatened (Bogoni et al. 2018) and it is crucial to measure the human activities influence on the environment and, ultimately, on these species’ distribution and density. Besides of collecting data on species occurrence and demographic trends its critical to collect data about anthropogenic pressures like plant exploitation, poaching intensity, invasive species dissemination and domestic animals’ presence on the environment. For this purpose, we suggest the creation of a local collected environmental quality indicator and aim to test whether this index correlates with already existing indices.

## METHODS

In this study we created an Anthropogenic Influence Index based on local collected data at the Atlantic Forest and compared it to traditional index such as the Global Human Influence Index, landscape ecology metrics (habitat cover, number of habitat fragments and matrix heterogeneity) and Human Development Index.

### Study area

We carried out our analyses in 24 study areas across the Atlantic Forest geographic distribution at the Brazilian territory (Instituto Brasileiro de Geografia e Estatística (IBGE 1992). The Atlantic Forest contains 15 ecoregions (Olson et al. 2001) and its remaining original cover (12 %) is highly fragmented allowing the availability of different landscape configurations to be studied (Ribeiro et al. 2009b). Our sampling sites were inside protected areas located between latitudes ranging from -10.7° to -29.5° and between longitudes ranging from -37.3° to - 53.9° (Fig. 1).

**Figure 1.**
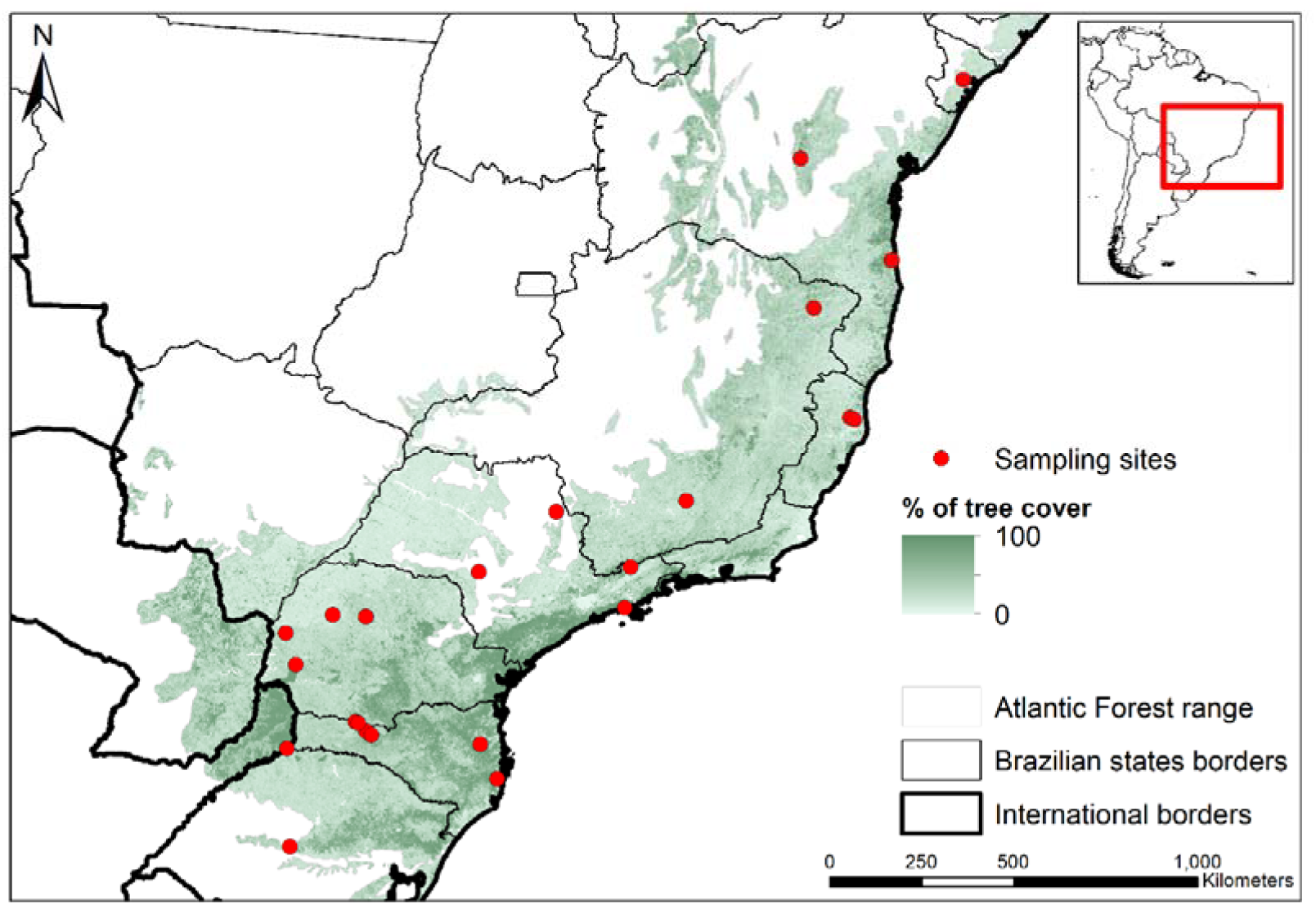
Sampling sites across the Atlantic Forest and percentage of tree cover (Hansen et al. 2003) showing its highly fragmented range.

### Anthropic influence index

The Anthropogenic Influence Index (AII) we created is constructed upon an assessment of five key factors in the field, encompassing: (i) poaching; (ii) exploitation of vegetation or invasion intensity (e.g., extraction of heart-of-palm, cultivation of cocoa and palms, *Pinus* invasion); as well as the presence of (iii) domestic ungulates (e.g cattle, sheep, lamb, etc), (iv) dogs, and (v) wild boars. Each of these factors were categorized into three categories: “absence,” “low frequency,” and “high frequency,” corresponding to scores of zero, one, or two, respectively. The AII is derived from the sum of the scores of each key factor and spans from zero to ten. Data regarding these factors were primarily gleaned from field expeditions focused on gathering biological samples (feces) for estimating cervid densities. Throughout these excursions, a multitude of direct and indirect indicators of the selected factors’ presence were noted while traversing the interior regions of protected areas. It was done without the use of previously established trails, and sampling efforts ranged from 486 m to 15,146 m. Furthermore, we also used to categorize the key factors informal discussions with protected area managers and researchers engaged in ongoing or past studies within these locales. Details of these categorizations is outlined in Table 1.

**Table 1.** Description and categorization of key factors used as components of the anthropogenic influence index (AII)

| Factors | Categories |  |  |
| --- | --- | --- | --- |
|  | 0 | 1 | 2 |
| Poaching | Probably nonexistent | Existing (Superficial reports) | Very frequent (Strong reports, local findings with bait stations, platforms, trails, traps, etc.) |
| Plant exploitation (heart-of-palm, cocoa, coffee, *Pinus*, Eucalyptus, oil palm, timber extraction) | Probably nonexistent | Existing (Superficial reports) | Very frequent (Strong reports, local evidence with presence of cut or planted specimens) |
| Domestic ungulates | Probably nonexistent | Existing (Superficial reports, present in surrounding properties) | Very frequent (Strong reports, local evidence with feces, sightings, etc.) |
| Domestic dogs | Probably nonexistent | Existing (Superficial reports, present and free ranging in surrounding properties) | Very frequent (Strong reports, local evidence with vocalizations, sightings, camera traps records etc.) |
| Wild boar | Probably nonexistent | Existing (Superficial reports) | Very frequent (Strong reports, local evidence with camera traps records, signs of activity, vocalizations, sightings, etc.) |

### Landscape metrics, human density, GHII and HDI

The landscape metrics were calculated within circular landscapes with a fixed radius of 2 km centered at the geographical coordinates of the central region where field incursions were conducted to obtain the AII. We utilized geoprocessing tools to delineate the various circular landscapes and extract the following variables from them: number of fragments, habitat coverage, and matrix heterogeneity. For this purpose, we based our analysis on a land use and cover map of 100m resolution obtained from Landsat images (Buchhorn et al. 2020) with data of year 2019. All forest cover categories from our base map of land use and cover were considered as habitats. We carried out these analyses using the R packages “landscapemetrics” (Hesselbarth et al. 2019) and “lanscapetools” (Sciaini et al. 2018).

We compared our AII with the GHII (Wildlife Conservation Society and Center for International Earth Science Information Network 2005) that was downloaded and had its pixels values extract to our study areas central geographical coordinates points. Additionally, we added to these points the human density (Center for International Earth Science Information Network - CIESIN - Columbia University 2018) and the subnational HDI extracted from data.worldbank.org for the same year of field data collection.

### Data Analysis

We organized our data set with each observation being a study area where we collected our AII and added to these observations the GHII, the human density, the habitat cover, the number of habitat fragments, the matrix heterogeneity and the subnational HDI spatially and temporally associated to it.

After normalizing the AII values we found in order to range from zero to one, we tested the linear correlation (Pearson) between AII and GHII, Human density, HDI and each landscape metrics. In addition, we carried out a multiple linear regression test where AII was the response variable and HDI, and the landscape metrics (number of patches, Matrix Shanon and habitat cover) were the predictive variables. The best model was validated by analyzing the plots of the residuals. All statistical analyses were carried out in R v. 4.2.3 (R Core Team 2019).

## RESULTS

We only found a significant correlation in our Person tests among the AII and the HDI (R = 0.51, df = 22, p = 0.01) with the other relations being nonsignificant (Fig. 2). Our multiple linear regression test indicate that the best model is AII ∽ HDI, where AII = - 6.9 + 9.7^*^HDI (F_1,22_ = 7.9, r^2^ = 0.23, P = 0.01) (Fig. 2).

**Figure 2.**
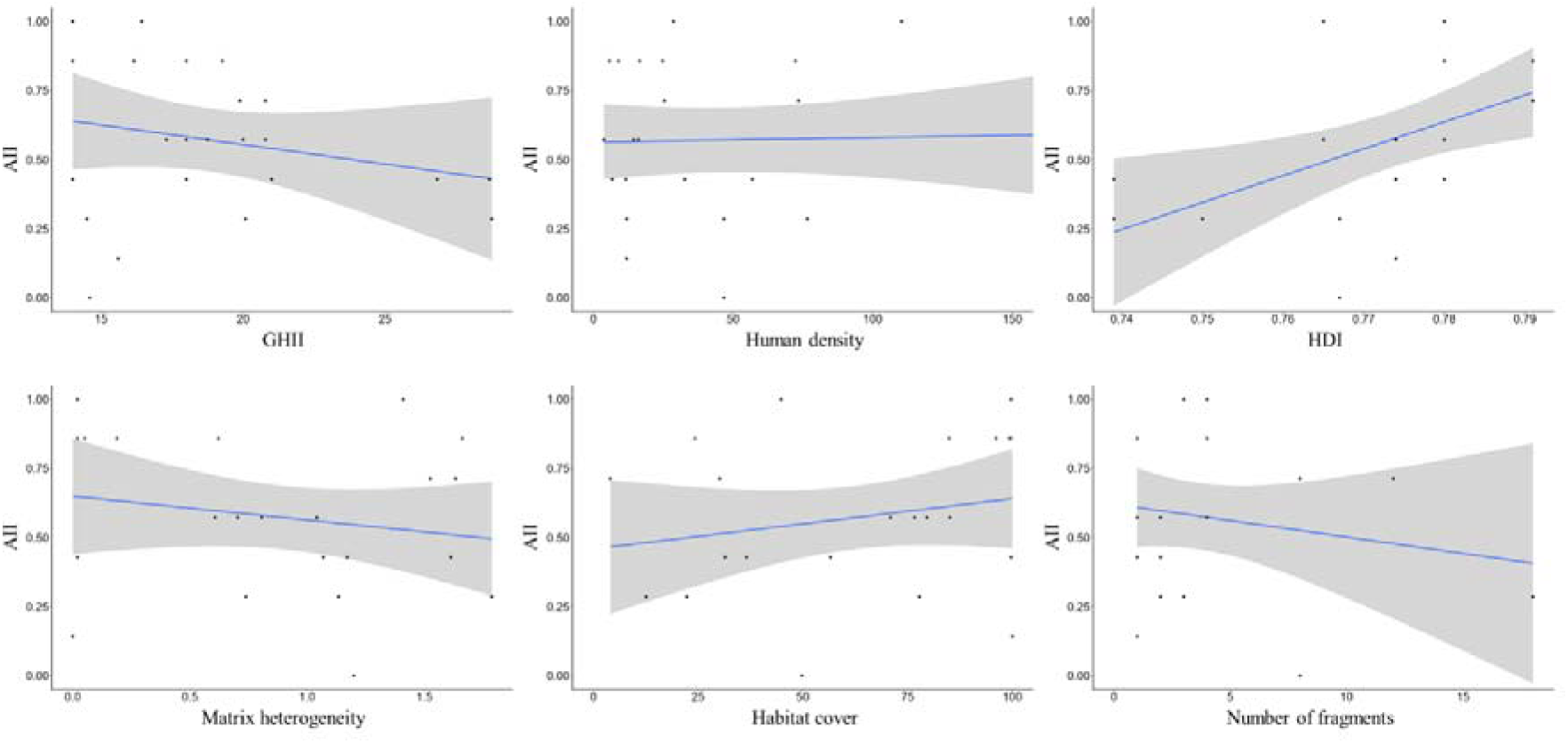
Pearson correlation test between the Anthropic Influence Index (AII) and the Global Human Influence Index (GHII)(R = -0.22, df = 22, p = 0.29), Human density (R = 0.04, df = 22, p = 0.84), Human Development Index (HDI)(R = 0.51, df = 22, p = 0.010*), matrix heterogeneity (R = -0.19, df = 22, p = 0.36), habitat cover (R = 0.21, df = 22, p = 0.31) and number of habitat fragments (R = -0.18, df = 21, p = 0.40).

## DISCUSSION

The absence of correlation between the AII and the GHII indicate that remote sensing-collected data, and inferences from it, may not always reflect local and specific anthropogenic impacts on the environment. In the case of the AII, it directly reflects the main potential impacts for medium and large mammals, apart from habitat loss, which is the most common data acquired in remote sensing-based studies such as land cover monitoring (Butti et al. 2022). In addition, the absence of correlation between the AII and the human population density may indicate that there is not a direct correlation between human population density and some anthropogenic impacts on the environment. This absence of correlation was found in other studies where the authors tested the correlation between human population density and land-based waste entering oceans (Schuyler et al. 2021) and intact mammal communities (Belote et al. 2020). In our case, this absence of correlation between AII and human population density indicates that anthropogenic impacts, especially those ones we detected calculating the AII, such as hunting pressure (Jerozolimski and Peres 2003), infection diseases from domestic animals (Portela Lobato et al. 2015), predation by domestic dogs (Cassano et al. 2014) and competition with exotic and invasive species (Doherty et al. 2017) may not be attributed to different levels of human density but to the presence of small settlements (Verlinden 1997) and rural properties (de Lima et al. 2020). Regarding the landscape metrics the absence of correlation between and the AII may indicate that the human impacts it detects are not directly associated with habitat availability, fragmentation levels and landscape heterogeneity. However, some studies have indicated that landscape metrics are associated with species richness (Palmeirim et al. 2019) and turnover (Beca et al. 2017).

The AII appears to serve as a robust indicator of environmental integrity, extending beyond considerations of habitat availability, fragmentation, and human population density. It brings to light the Empty Forest paradigm, a hypothesis initially proposed by (Redford 1992), which suggests that forests seemingly intact on the surface may suffer from defaunation due to factors such as overhunting, habitat loss, and competition for resources with livestock (Ripple et al. 2015). To investigate the empty forest hypothesis, the AII may be an interesting environmental quality indicator if replicated in other studies. However, it is important to note that the AII relies on field-collected data, necessitating labor-intensive expeditions with significant human and financial costs compared to remote sensing-based methods. Therefore, careful consideration is warranted when choosing between more informative yet expensive environmental quality indicators and less informative but more affordable methods. In certain contexts, only detailed, locally collected indicators like the AII can offer insights into conservation challenges (László et al. 2014; Frasconi Wendt et al. 2021).

Scale is a fundamental concept in ecology and environmental sciences (Hernández 2020). Depending on the scale, certain natural phenomena or anthropogenic impacts on the environment may go undetected if they exhibit local particularities in studies conducted at broad scales. Conversely, general patterns may be overlooked in studies at narrow scales, which are more susceptible to biases caused by lack of detail. Clear examples of issues related to scale come from landscape ecology studies that have identified that certain phenomena may only be detected or modeled at particular scales, considering both resolution and extent (Mayer and Cameron 2003).

The significant correlation between AII and HDI draws attention to the direct relationship between income level, level of education, and life expectancy, the components of HDI, and the incidence of environmental impacts. Intuitively, we would expect the opposite: an inverse relationship between socioeconomic development and environmental impact. However, it has been reported that deforestation rates increase with the rise of socioeconomic indicators (Michinaka et al. 2020). In this sense, it is argued that the relationship between socioeconomic indicators and environmental issues behaves like a Kuznets curve (Stern 2004), where there is an initial phase where the relationship is positive, as we have detected, and in a second phase it becomes negative (Cropper et al. 1999). Regarding local communities, economic inequality is shown to be more associated with drivers of illegal hunting than poverty in general (Lunstrum and Givá 2020).

Reflecting on human development involves addressing various sources of deprivation and oppression worldwide. Expanding freedom is both the goal and means to achieve development (Sen 1999). While the Human Development Index (HDI) measures health, education, and income, it lacks indicators for other crucial aspects like participation, democracy, tolerance, equity, and sustainability. Complementing the HDI with poverty and environmental metrics is essential. Developing adjusted versions such as the Inequality-adjusted HDI (IDHAD) and Planetary Pressures-adjusted HDI (IDHP) would enhance its comprehensiveness (PNUD 2021).

Analyzing human influence at the local level, as proposed by our Anthropogenic Influence Index, allows for the specification and detailed collection of socio-environmental data. This approach considers cultural and biological distinctions to create or improve criteria tailored to managing conservation goals effectively. Nonetheless, refining and adapting the Anthropogenic Influence Index necessitates further research experiments.

Finally, we emphasize that while the AII appears to better capture local nuances of environmental impacts, particularly those significant for medium and large mammals, compared to other indicators such as GHII, human density, and landscape metrics, further investigation is necessary to understand the effects of different levels of AII on mammal occurrences, occupancy dynamics, and population densities. Moreover, it would be intriguing to explore how ecological relationships such as predation and competition respond to habitats with heterogeneous AII levels.

## ACKLODEDGEMENTS

We are indebted to two anonymous reviewers who reviewed our manuscript carefully and in-depth, considerably improving its quality. This study was funded by the Sao Paulo Research Foundation (FAPESP) (2015/25742-5), the National Council for Scientific and Technological Development (CNPq) (314038/2021-3, 150222/2022-0).

